# Myocardial matrix material supports a proliferative microenvironment for cardiomyocytes

**DOI:** 10.1101/2020.07.31.231845

**Authors:** Raymond M. Wang, Paola Cattaneo, Nuno Camboa, Rebecca Braden, Colin Luo, Sylvia Evans, Karen L. Christman

## Abstract

Novel therapeutics have sought to stimulate the endogenous repair mechanisms in the mammalian myocardium as the native regenerative potential of the adult cardiac tissue is limited. In particular, a myocardial matrix derived injectable hydrogel has shown efficacy and safety in various animal myocardial infarction (MI) including evidence of increased myocardium. In this study, investigation on the properties of this myocardial matrix material demonstrated its native capability as an effective reactive oxygen species (ROS) scavenger that can protect against oxidative stress and maintain cardiomyocyte proliferation *in vitro. In vivo* assessment of of myocardial matrix hydrogel treatment post-MI demonstrated increased thymidine analog uptake in cardiomyocytes compared to saline controls along with co-staining with cell cycle progression marker, phospho-histone H3. Overall, this study provides further evidence that properties of the myocardial matrix hydrogel promote an environment supportive of cardiomyocytes undergoing cell cycle progression.

## Introduction

Ischemic heart disease contributing to myocardial infarction (MI) and subsequent heart failure (HF) is a leading cause of death in the western world and worldwide accounting for 15.5% mortality in 2015^1, 2^. Although medical response to acute MI has led to reduced death from these incidents, the limited regenerative potential of the adult mammalian myocardium leads to continued cardiomyocyte apoptosis, negative tissue remodeling, and eventually heart failure. Current treatments for end-stage heart failure include heart transplant and left ventricular assist device, however, limitations on available healthy hearts for transplant^3^, strain on medical resources, and effects on patient quality of life have prompted the need for improved treatments. Thus, extensive research has been focused on developing novel treatment methods to restore the damaged cardiac muscle.

Traditionally, the mammalian heart was considered to have no significant ability to regenerate damaged cardiac muscle tissue. However, studies in both mice and porcine cardiac injury models have determined that the mammalian heart during the fetal and neonatal stages immediately post-gestation can regenerate large regions of infarcted tissue without significant scar tissue formation^4-7^. These results demonstrate that the mammalian heart can utilize mechanisms of significant myocardial regeneration from pre-existing cardiomyocytes during early life stages that is rapidly lost with aging. Research to determine why adult cardiomyocytes lose regenerative capability has suggested cell cycle progression is arrested shortly after birth due to changes in microenvironmental cues such as localized hypoxia reducing oxidative stress^8, 9^, and extracellular matrix (ECM) composition^10-12^. Based on these principles, hypoxia fate mapping techniques have found incidents of cardiomyocytes, termed Hif-1α cardiomyocytes, with proliferative characteristics in the adult myocardium in localized hypoxic microenvironments. These cardiomyocytes maintain immature characteristics and turnover rates similar to previously determined rates in human hearts suggesting their contribution^13^. Furthermore, inducing systemic hypoxia^8^ and increasing or decreasing sources of oxidative stress^14^ has been shown to influence the numbers of these cardiomyocytes *in vitro* and *in vivo*, thus, suggesting these potential routes for alternative therapies to stimulate cardiomyocyte renewal.

Amongst these alternative therapies, a decellularized myocardial matrix hydrogel derived from left ventricular porcine tissue has been demonstrated as a post-MI therapy leading to mitigation of negative left ventricular remodeling and loss of function in small and large animal models compared to controls^15, 16^, leading to evaluation in a Phase I clinical trial^17^. Notably, analysis of the matrix treated tissue provided evidence suggesting some promotion of mechanisms to restore the cardiomyocyte population. In the MI rat model, larger clusters of cardiomyocytes within the infarct region in myocardial matrix versus saline groups were observed after 4 weeks post-injection^16^. Transcriptomic microarray analysis of the infarcted rat heart tissue demonstrated that the myocardial matrix treatment versus saline promoted genes related to reduced cardiomyocytes apoptosis, shifted cardiac metabolism, and greater cardiac development^18^. Finally, in a large animal porcine MI model, an increased endocardial muscle layer was observed with matrix delivery 2 weeks post-MI compared to saline and non-treated controls when evaluated at 3 month post-injection. In contrast, saline injected and non-treated controls contained minimal remaining cardiac muscle^19^.

Based on literary support of the influence of oxidative stress on cardiomyocyte cell cycle progression and observations of influence on cardiac tissue repair from myocardial matrix treatment, the goal of this study was to investigate whether the properties of the myocardial matrix hydrogel material provided a microenvironment conducive to support cardiomyocyte cell cycle progression, and evaluate whether this was a contributing mechanism to the observed therapeutic efficacy.

## Methods and Materials

### Preparation of Myocardial Matrix Material for *in vitro* and *in vivo* application

Injectable aliquots of myocardial matrix material were created and assessed for material characteristics based on previously described protocols^20^. In brief, fresh porcine hearts were excised from adult pigs and the left ventricular tissue was isolated. Major vessels and fascia were removed and remaining muscle tissue was cut into less than 5 mm sized pieces. Tissue was then decellularized in 1% sodium dodecyl sulfate solution in 1x PBS under agitation for 4-5 days until tissue appeared completely white with an additional day of rinsing in water and repeated water rinsing with manual mixing. Decellularized tissue was lyophilized and milled from at least three hearts into a fine powder. Milled myocardial matrix was partially digested with a tenth mass of pepsin relative to matrix material in 0.1 M HCl at room temperature for 48 hours. Solution was neutralized with 1 M NaOH to a pH of 7.4. Neutralized myocardial matrix was aliquoted, frozen in a -80°C freezer, lyophilized, and stored with desiccate for long term storage at -80°C until needed for use.

### Material Characterization

Capability of myocardial matrix material as a reactive oxygen species (ROS) scavenger was investigated by reconstituting myocardial matrix in a microcentrifuge tube with 100 µL of 1x PBS at a 6mg/mL concentration and forming hydrogels overnight. Collagen gels at 2.5 mg/mL concentration were used for controls for similar porosity and mechanical strength to matrix gels^21^. 150 µL of 166µM H_2_O_2_ peroxide solution diluted in PBS was applied to each gel (100 µM final concentration). A solution only control in 1x PBS to assess degradation of the hydrogen peroxide over time was also made. Samples were incubated at 37°C on a shaker plate set to 120 rpm and hydrogen peroxide content was determined by Pierce™ Quantitative Peroxide Assay Kit. Similar procedural set-up was applied to matrix and collagen material in solution along with pre-milled myocardial matrix scaffold material and milled material incubated with hydrogen peroxide to determine influence of material processing on scavenger activity. Total antioxidant activity and thiol content of myocardial matrix material and collagen controls was measured by Cayman Chemical’s Antioxidant Assay kit and Cayman Chemical® Thiol Detection Assay Kit, respectively. Results were normalized to mass of material for comparison of antioxidant activity and thiol content.

### Neonatal Cardiomyocyte Isolation and Hydrogel Encapsulation

P1 neonatal rat cardiomyocytes were isolated from Sprague Dawley rat pups with a Neonatal Cardiomyocyte Isolation System (Worthington Enzyme). Following manufacturer instructions collecting cardiac cell suspension from neonatal myocardial rat tissue, the cardiac cell suspension was pre-plated for 2 hours and the cells in suspension were collected to enrich for cardiomyocytes. For assessment of cell proliferation, cells were pre-labeled by CellTrace™ Far Red (ThermoFisher Scientific) before encapsulation while other cellular experiments proceeded with unlabeled cells. Encapsulation of cells was performed with some protocol modifications to previously described protocols for improving viability for this specific cell type ^22^. In brief, solely neutralized myocardial matrix in lyophilized aliquots were resuspended to a 6mg/mL concentration with culture media consisting of 1:1 DMEM/F12 solution (Gibco) supplemented with 10% fetal bovine serum (Gibco) and 1% penicillin streptomycin (Gibco). Cells for encapsulation were pelleted by 50 rcf centrifugation at 4°C and resuspended with the matrix culture media mixture. 25 µL gels were formed containing 300,000 cells per gel by pipetting the mixture as a collected droplet in the well center of 24 well plates and incubating in a 37°C incubator for 1 hour. Following the 1 hour incubation, gel formation was confirmed by tilting the well plate vertically and observing maintained gel structure before adding 600 µL of culture media to each well. Encapsulated cells were cultured at 37°C and 5°C CO_2_ with media changed every 2-3 days.

### Flow Cytometry Analysis

CellTrace™ prelabeled cells from encapsulations were removed from the well plate surface by a sterile spatula and at least three gels were batched per sample into an enzymatic digestion solution consisting of 1:1 solution of HBSS (calcium and magnesium supplemented) and 1% bovine serum albumin in PBS with 1 µM HEPES (Gibco), 300 U/mL collagenase type IV (Worthington Biochemical), and 10 U/mL DNase I (Sigma-Aldrich). Gels in enzymatic digestion solution were incubated at 37°C under mechanical agitation at 600 rpm on a thermomixer (Benchmark Scientific) for 20 minutes. Solutions were then kept in ice and FACS buffer consisting of 2% fetal bovine serum and 1mM EDTA in DPBS lacking calcium and magnesium to inactivate enzyme activity. Cells were centrifuged at 50 rcf centrifugation at 4°C and resuspended in HBSS. Cell suspension was stained with LIVE/DEAD™ Fixable Aqua (ThermoFisher Scientific) for 15 minutes at 4°C and excess dye was quenched with FACS buffer. Cells were fixed for 10 minutes and permeabilized with BD Cytofix/Cytoperm™ solution, respectively. PCM-1 antibody (1:400, Sigma-Aldrich) and concentration matched rabbit IgG isotype control (Novus Biologicals) were incubated with cells for 12-18 hours at 4°C in BD Cytoperm™ solution. Secondary staining with donkey anti-rabbit PE (1:1500, Biolegend) was applied for 30 minutes at 4°C and stained cells were resuspended in FACS buffer before flow analysis on a BD FACSCanto™ II (BD Biosciences). Gating and flow data were processed on FlowJo (FlowJo LLC).

### Metabolic and Gene Expression Analysis

Encapsulated cells were assessed for redox related metabolic activity based on resazurin oxidation-reduction indicator in viability alamarBlue™ (ThermoFisher Scientific) assay. Encapsulated cells in matrix or collagen control gels were cultured in culture media alone and under oxidative stress by supplementing 2.5 mM H_2_O_2_ mixed into the culture media for 4 hours. Following 4 hour incubation, gels were removed from the well plate surface by a sterile spatula and at least three gels were batched per sample for RNA isolation. RNA was isolated by RNEasy kit (Qiagen, Germantown, MD) along with an on-column DNase digestion step (Qiagen) to extract RNA with minimal genomic DNA contamination. Superscript III Reverse Transcriptase kit (Applied Biosystems, Foster City, MA) was used to synthesize cDNA. Then, SYBR Green PCR Master Mix (Applied Biosystems) was used with forward and reverse primers at a final concentration of 200 nM.

Gene expression of cell cycle regulators p19ARF and p16 that are upregulated in response to oxidative stress leading to cell cycle arrest and apoptosis were assessed. Primers sequences were as follows: rat p19ARF (F: 5’-GGTTTTCTTGGTGCAGTTCCTG-3’, R: 5’-GATCCTCTCTGGCCTCAACAC-3’), rat p16 (F: 5’-GATCCAGGTCATGATGATGGG-3’, R: 5’-ATCAATCTCCAGTGGCAGCG-3’) and 18s rRNA (F; 5’-GGATCCATTGGAGGGCAAGT-3’, R: 5’-CCCAAGATCCAACTACGAGCTT-3’). Samples were run in technical duplicates along with negative controls without template cDNA to confirm lack of contamination in PCR reagents. PCR reactions were run on a CFX95™ Real-Time System (Biorad, Hercules, CA) with the following thermal cycler settings: 30s at 50**°**C, 2 min at 95**°**C, 40 cycles of 10s at 95**°**C, and 30 secs at 63**°**C based on pre-determined optimal primer efficiency amplification temperature. After completing 40 cycles of PCR amplification, automated melting curve analysis, consisting of increasing the thermal cycler temperature from 50**°**C to 95**°**C at 5**°**C increments lasting 5s each, was used to confirm formation of a singular PCR amplicon for each primer set. Bio-Rad CFX Manager™ 3.0 (Biorad) was used for determining cycle threshold (ct) values from recorded SYBR green signal. Fold change was then determined by 2^-(Gene 1 – 18s)^ and normalized to fold change of respective material sample without H_2_O_2_ doped into the media.

### *In Vivo* Delivery of Thymidine Analogs Following Myocardial Matrix Hydrogel Delivery Post-myocardial infarction

All experiments in this study were performed in accordance with the guidelines established by the committee on Animal Research at the University of California, San Diego, and the American Association for Accreditation of Laboratory Animal Care. Two groups of Sprague Dawley rats (n = 8 per group) underwent a previously described ischemia-reperfusion procedure^16^ accessing the heart by left thoracotomy for 35 minute temporary occlusion of the left coronary artery to induce MI. After 1 week post-MI, animals were randomized for myocardial matrix or saline injection. Myocardial matrix aliquots that were salt reconstituted were resuspended with sterile water into a homogeneous suspension before injection. The heart was viewed through an excision in the diaphragm to deliver a 75 µL direct intramyocardial injection into the infarct area^16, 18^. Following the injection procedure, Alzet® osmotic pumps were implanted subcutaneously in the dorsal region as previously described^23^ providing continuous delivery of 20 mg/kg/day of 5-Ethynyl-2’deoxyuridine (EdU) for 1.5 weeks. Afterwards, the initial pump was replaced with a second pump containing an alternative thymidine analog bromodeoxyuridine (BrdU) for an additional 1.5 weeks. To account for expected delay of thymidine analog getting to the heart based on subcutaneous delivery, intraperitoneal injections of each respective thymidine analog at a concentration of 20 mg/kg was performed on the day preceding, immediately following, and the day after pump implantation. At day 29 post-MI, rats were euthanized by 300 µL lethal dose of sodium pentobarbital delivered by intraperitoneal injection, excision of the heart, and fresh freezing the tissue in OCT for sectioning.

### Histology and Immunohistochemistry

Fresh frozen tissue in OCT was cryosectioned to obtain transverse sections of the heart. Cryosections from 12-16 different evenly spaced locations were used for all immunohistochemistry. For infarct size analysis, hematoxylin and eosin staining and quantification of percent infarct area by ratio of infarct area to left ventricular and septal area was used to assess for outliers that should be excluded from subsequent analysis.

For immunohistochemical staining, slides were fixed with acetone or 4% paraformaldehyde for 5 minutes at -20°C and 10 minutes at room temperature, respectively, and permeabilized with 0.5% Triton-X in PBS for 10 minutes. EdU incorporation was stained by Click-iT™ Plus EdU Cell Proliferation Kit for Imaging, Alexa Fluor™ 647 dye (ThermoFisher Scientific) or Click-&-Go™ Plus 647 Imaging Kit (Click Chemistry Tools, Scottsdale, AZ) preceding application of antibody staining solutions. For antibody staining, sections were blocked with a buffered solution containing 5-10% donkey serum, 1% bovine serum albumin, and 0.1% Triton-X 100 based on the optimized antibody protocol.

The following primary antibodies were incubated for 1 hour at room temperature or for 12-18 hours at 4**°**C for the following markers: rabbit PCM-1 (1:100 dilution, Sigma-Aldrich, HPA023370-100UL), mouse PCM-1 (1:200, Santa Cruz, sc-398365), α-actinin (1:800, Sigma-Aldrich, A7811-100UL), goat PDGFR-α (1:200, Novus Biologicals, AF1062), goat vimentin (1:200, Santa Cruz, sc-7557), Isolectin Griffonia Fluorescein (1:150, Vector Laboratories, FL-1201), mouse smooth muscle cells (1:800, α-SMA, M0851), mouse CD68 (Biorad, 1:200, MCA341GA), mouse CD45R (1:200, BD Pharmingen, 554879), BrdU (1:100, ThermoFisher Scientific, B35128), and rabbit pHH3 (Cell Signaling, 9701S).

The following secondary antibodies were incubated for 30-45 minutes at room temperature: anti-rat Alexa Fluor 568 (1:500 dilution), anti-rabbit Alexa Fluor 488 (1:800 dilution), anti-mouse Alexa Fluor 488 (1:800 dilution), anti-rat Alexa Fluor 488 (1:500 dilution), anti-rabbit Alexa Fluor 568 (1:500 dilution), anti-rabbit Alexa Fluor 647 (1:500 dilution), and Hochest 33342 (Sigma-Aldrich, H3570) was applied as a nuclei counterstain. Slides were rinsed and coverslipped with Fluoromount™ Aqueous Mounting Medium (Sigma-Aldrich). Bright field images were taken a Leica Aperio ScanScope® CS2 and fluorescent images with a Leica Ariol® system (Leica). Selection of region of interest around the infarct and confirmation of positive colocalization was drawn and visualized, respectively, in Aperio ImageScope software (Leica). Automated expansion around the infarct at 250 µm intervals for designating a borderzone region of interest for analysis and automated analysis of costaining was performed by custom MATLAB scripts (Mathworks, Natick, MA).

### Statistical Analysis

All data and plots are presented as mean ± SD. Significance was determined with a one-way ANOVA using a Tukey post-hoc test and an unpaired student’s *t*-test based on number of sample groups with significance designated at p<0.05.

## Results

### Decellularized Myocardial Matrix Function as a ROS Scavenger

Matrix interaction with reactive oxygen species (ROS) was examined based on studies demonstrating that reduced oxidative stress promotes proliferative characteristics in cardiomyocytes during the neonatal and adult stages^9, 24, 25^. Concentration of incubated H_2_O_2_ with myocardial matrix hydrogels versus collagen controls was measured over 5 days. These measurements determined a significantly continuing decrease of H_2_O_2_ concentration for the myocardial matrix versus both PBS and collagen controls. Collagen also had a significant decrease to PBS, though the difference plateaued after a day, while H_2_O_2_ in PBS maintained relatively consistent (Figure 1A). Assessment of this response for decellularized materials in various forms determined that this activity is related to spatial access to sites of ROS scavenger activity in the decellularized material, with less consistent reduced H_2_O_2_ for whole decellularized ECM versus ECM powder or hydrogel form (Figure S1). Further confirmation and relative strength as an antioxidant were assessed with a Cayman Chemical’s Antioxidant Assay kit based on comparison with an antioxidant standard, Trolox. In comparison to Trolox, an equivalent scavenger activity of 4.00 ± 0.359 mM Trolox/mg myocardial matrix was determined compared to 1.02 ± 0.840 mM Trolox/mg collagen (Figure 1B). Since general ROS scavenger activity of protein mixtures is commonly linked to specific chemical groups such as thiol groups found in available free cysteine amino acids, we utilized a Cayman Chemical® Thiol Detection Assay Kit to determine thiol content based on a glutathione standard curve, which similarly showed around a fourfold greater thiol content compared to collagen of 75.8 ± 13.2 nmol/mg myocardial matrix versus 22.0 ± 10.2 nmol/mg collagen (Figure 1C).

**Figure 1:**
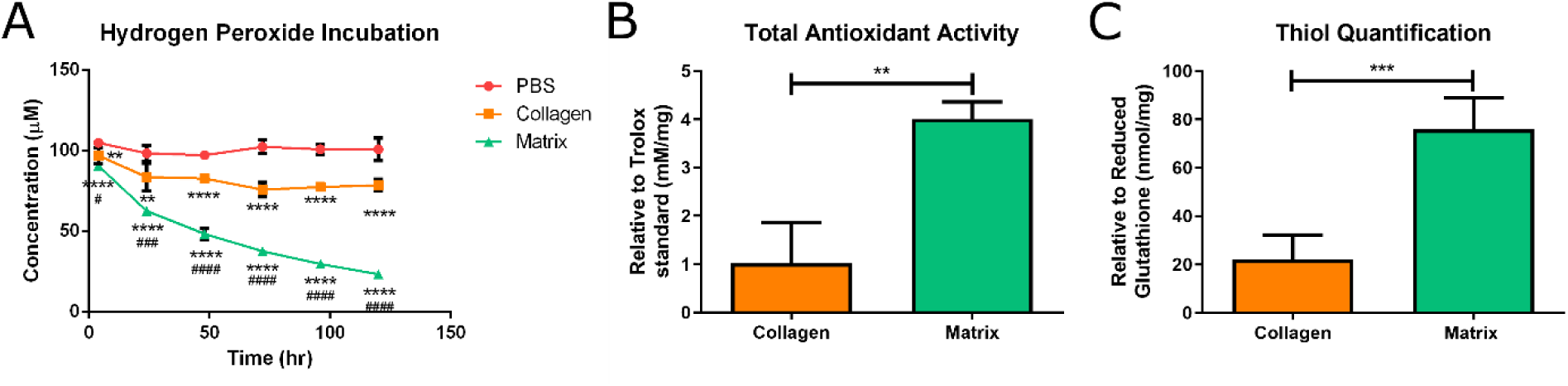
ROS scavenger capability and comparison to antioxidants of myocardial matrix hydrogel. (A) 150 µL of 166.7 µM hydrogen peroxide diluted in PBS solution was applied to a solution only control (red), and 100 µL collagen (orange) and myocardial matrix (green) hydrogels (n = 5 each). Measurements at 4, 24, 48, 72, 96, and 120 hours of the hydrogen peroxide in the supernatant were measured showing a continuing decrease in concentration for myocardial matrix versus PBS and collagen controls. (B) Total antioxidant activity determined relative a Trolox antioxidant standard for myocardial matrix versus collagen in solution per mg of material (n = 4) (C) Thiol content was determined compared to glutathione standard in myocardial matrix versus collagen per mg of material (n = 4 each) (* is significance to PBS, # is significance to collagen gel, \**p*< 0.05, \*\**p*< 0.01, \*\*\**p*< 0.001, \*\*\*\**p*< 0.0001).

### Myocardial Matrix Reduces Effects of Oxidative Stress

To determine whether this ROS scavenging environment lead to changes in cardiomyocyte phenotype, encapsulated neonatal cardiomyocytes after 2-3 days were treated with 2.5mM H_2_O_2_ in culture media for 4 hours. Change in cell viability related to redox metabolic activity was measured with AlamarBlue assay showing a significantly greater change compared to cells in collagen gels (Figure 2A). mRNA was then isolated from encapsulated cells and expression of cell stress markers that inhibit of cell cycle progression during oxidative stress, p19arf and p16, were measured compared to 18s rRNA housekeeping gene. Difference in cycle number values relative to respective control gels in 10% fetal bovine serum supplemented DMEM/F12 media determined a significantly greater expression of these stress markers in collagen culture gels compared to matrix suggesting a reduction in activation of oxidative stress pathways (Figure 2B).

**Figure 2:**
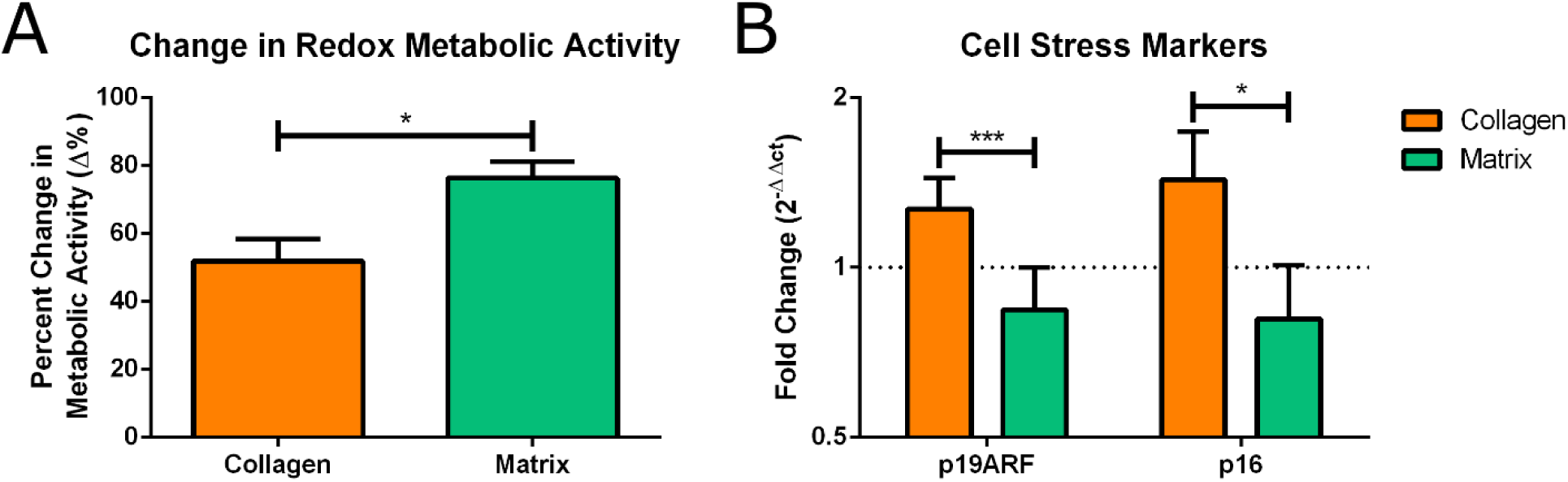
Cellular metabolism and stress markers of encapsulated cardiomyocytes in response to oxidative stress. 2.5mM hydrogen peroxide in 500µL of basal DMEM/F12 media was applied to 250,000 encapsulated neonatal cardiomyocytes in 25 µL collagen (orange) hydrogels and myocardial matrix (green) hydrogels for 4 hours. (A) Change in metabolic activity was determined with AlamarBlue assay, which is known to respond to alterations in redox metabolic activity (n = 4 each). (B) Assessment into expression of oxidative cell stress markers that inhibit cell cycle progression, p19ARF and p16, were determined relative to housekeeping gene, GAPDH (n = 4 each). (* is significance to PBS, # is significance to collagen gel, \**p*< 0.05, \*\**p*< 0.01, \*\*\**p*< 0.001, \*\*\*\**p*< 0.0001).

### Increased proliferation of *in vitro* encapsulated neonatal cardiomyocytes

Encapsulated P1 neonatal cardiomyocytes were then assessed for differences in cell turnover in myocardial matrix versus collagen hydrogel controls. Utilizing pre-labeling with CellTrace™, a proliferation assay was performed based on serial dye dilution with each progeny generation to confirm whether cardiomyocytes underwent increased incidents of cytokinesis. Gating for analysis of progeny by dye dilution on a flow cytometer was confirmed by pre-labeling of cardiac cell populations in 2D culture on tissue culture plastic. Samples were collected during various steps of the cell processing and after days of culture (Figure 3A).

**Figure 3:**
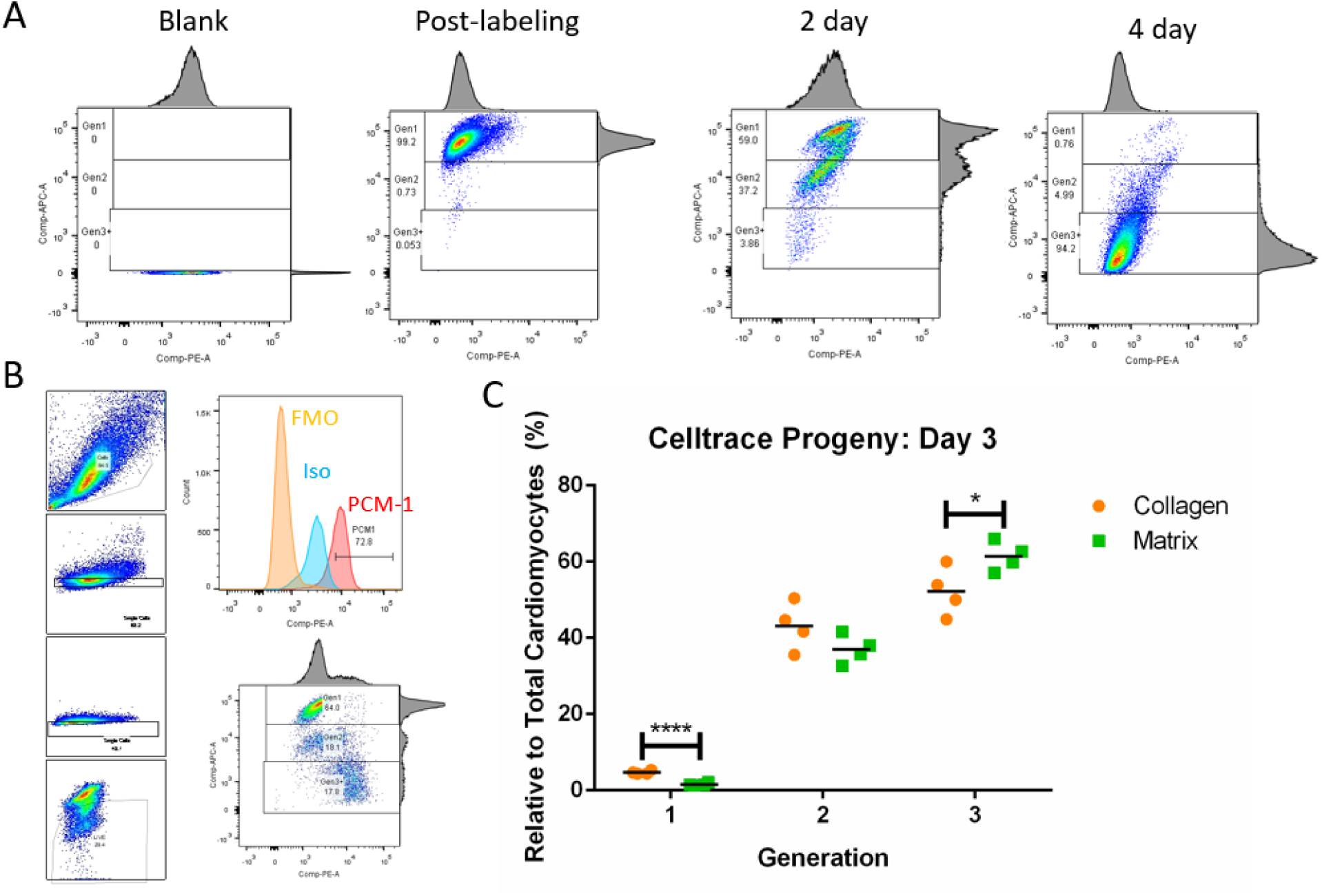
Progeny tracking of encapsulated cardiomyocytes. (A) Samples for optimizing gating for proliferative analysis based on CellTrace™ FarRed. Gating was based on non-label cells, immediately labeled cells, pre-labeled cells following 2 days of culture, and 4 days of culture demonstrating distinct clustered dye dilution representing increased generation of progeny. (B) Representation gating scheme for distinguishing live singlet cells based on forward and side scatter and LIVE/DEAD™ Aqua signal. PCM-1 gating with fluorescence minus one and isotype control peaks for distinguishing the cardiomyocyte positive peak and distribution of progeny. (C) Percentage of cardiomyocytes over two progeny generations after 3 days of culture encapsulated in collagen or myocardial matrix hydrogels. \**p*< 0.05, \*\*\*\**p*< 0.0001).

For specific determination of live cardiomyocytes, staining for LIVE/DEAD™ Aqua and PCM-1, cardiomyocyte nuclei marker, was utilized. PCM-1 has been used in mice and human cardiomyocyte studies, but has not been extensively validated for cardiomyocyte nuclei staining in rat tissue^26^. For validation, PCM-1 was co-stained in rat heart tissue sections with another cardiomyocyte marker (α-actinin) along with markers for fibroblasts (PDGFR-α, vimentin), endothelial cells (PECAM-1), smooth muscle cells (α-SMA), immune cells (CD68, CD45). Results showed overlapping co-stain with α-actinin and no co-labeling with non-cardiomyocyte markers confirming PCM-1 is a cardiomyocyte nuclei specific marker^27^ (Figure S2A-H). Further validation was performed with an alternative cardiomyocyte nuclei marker, Nkx2.5, showing colocalization of the two stains supporting specific labeling with PCM-1 (Figure S3).

Gating for PCM-1 by flow cytometry was validated by fluorescent minus one and IgG isotype control (Figure 3B). Analysis of encapsulated cardiomyocytes with these optimized gates determined a significant decrease in the original generation 1 cardiomyocyte population and a significant increase in cardiomyocyte progeny of the third generation relative to progeny distribution in collagen controls demonstrating increased proliferation of neonatal cardiomyocytes in the matrix material (Figure 3C).

### *In vivo* incorporation of thymidine analog in cardiomyocyte nuclei

For analysis, hematoxylin and eosin staining and quantification of percent infarct area was performed to assess for outliers that should be excluded from subsequent analysis. From this assessment, two hearts (one from each group) were immediately excluded due to lack of visible infarct. The infarct size in the remaining hearts (n = 7 per group) was then quantified, demonstrating there was no significant difference between groups (Figure S4). Thus, these hearts were carried over for subsequent analysis.

For analysis of thymidine analog uptake, co-staining of EdU or BrdU with PCM-1 was performed in the infarct and borderzone region. For consistency, the borderzone was identified by highlighting the infarct region in stained sections and expanding outwards from this highlighted region by 250µm up to 750µm (Figure S5A). Analysis of these two regions identified a significantly increased EdU^+^PCM-1^+^ density in the borderzone but not infarct region of matrix treated hearts versus saline treated hearts (Figure 4). Similarly, staining for BrdU^+^PCM-1^+^ cardiomyocytes were mainly identified in the borderzone region (Figure 4D), however, no significant differences were found from BrdU^+^PCM-1^+^ analysis in the borderzone region (Figure 4E) and no BrdU^+^PCM-1^+^ cardiomyocytes were found in the infarct region (not shown).

**Figure 4:**
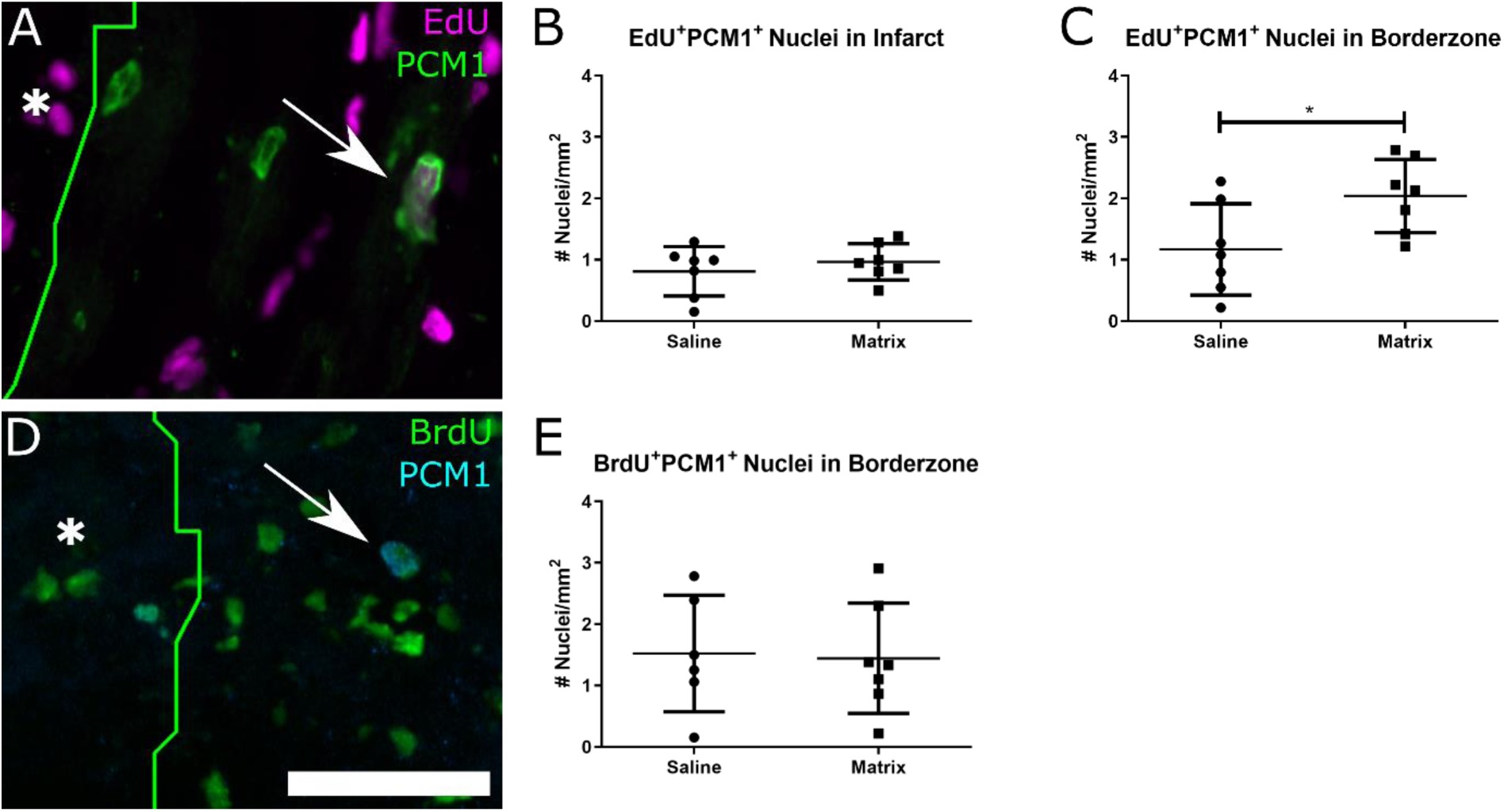
Thymidine analog uptake in cardiomyocyte nuclei. (A) Image of nuclei co-stain of thymidine analog EdU (purple) and cardiomyocyte nuclei specific marker PCM-1 (green) labeled by a white arrow, and white asterisk indicating infarct region. Density EdU^+^PCM-1^+^ nuclei in the (B) infarct. (C) infarct borderzone. (D) Image of nuclei co-stain of thymidine analog BrdU (green) and cardiomyocyte nuclei specific marker PCM-1 (blue) labeled by a white arrow, and white asterisk indicating infarct region. (E) Density BrdU^+^PCM-1^+^ nuclei in the infarct borderzone. (*p < 0.05, unpaired Student’s t-test)

### Staining for Cell Cycle Progression Markers

To provide further evidence that cardiomyocytes were undergoing cell cycle progression, PCM-1 was co-stained with cell cycle marker phospho-histone H3 (pHH3) from hearts isolated at day 7 post-injection based on increased positive thymidine analog co-staining with PCM-1 within this timespan and previous microarray data showing increased expression of genes related to cardiac development^28^. From these stains, we observed low incidents of co-staining with PCM1 with pHH3 (Figure 5A, B), supporting that incidents of thymidine analog uptake involved further progression into the cell cycle past S phase and were not just indications of DNA repair responses.

**Figure 5:**
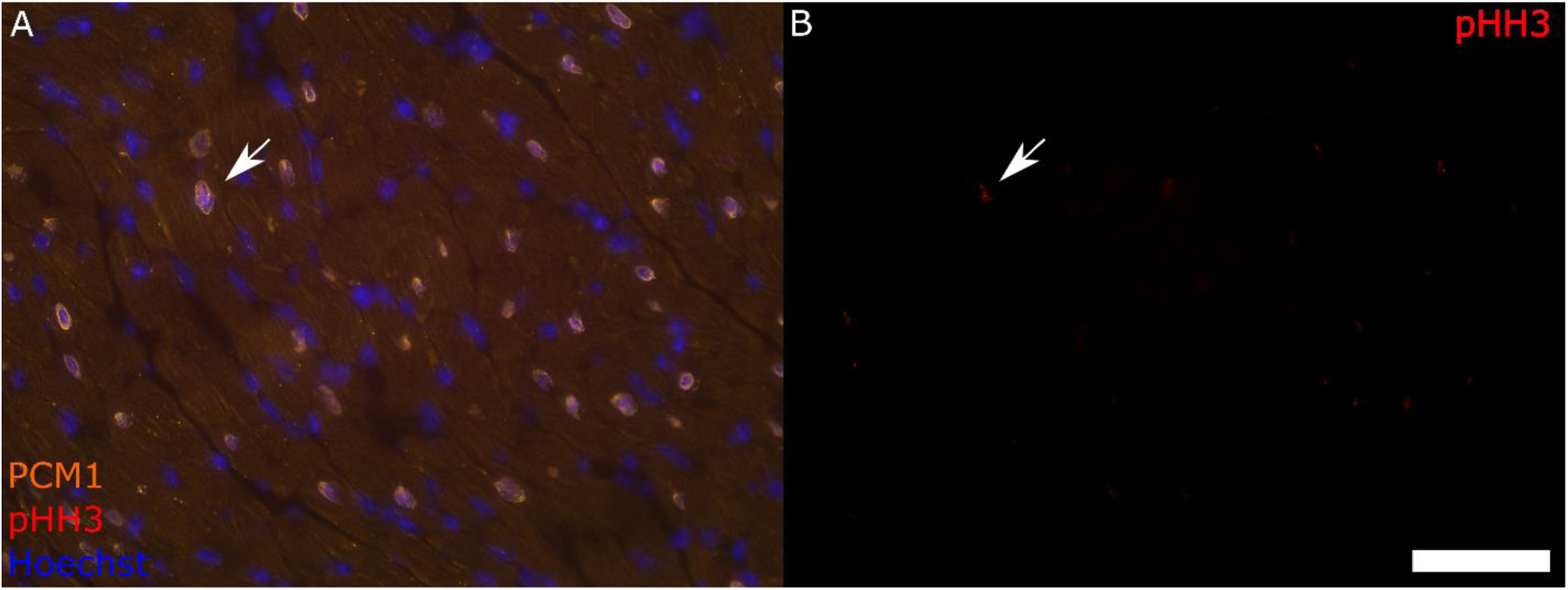
Staining of phosphor-histone H3 with cardiomyocyte nuclei. (A) Image of cardiomyocyte nuclei co-stain based PCM1 (orange) with pHH3 (red) with Hoechst counterstain (blue) with co-staining labeled by a white arrow. Scale bar of 100 µm.

## Discussion

As cardiac injury from MI is known to lead to sudden and continuous cardiomyocyte death, various therapies have attempted to renew the lost cardiomyocyte population as a therapeutic strategy. Although there has been various hypotheses and controversies on the potential and source of new cardiomyocytes to form in the mammalian adult myocardium, the following hypotheses are currently dominate in the field: 1) the mammalian heart up to shortly post-gestation has significant proliferative capability^4-7^, 2) mammalian cardiomyocyte renewal occurs from pre-existing cardiomyocytes similar to mechanisms seen in common non-mammalian animal models for studying cardiac repair such as zebrafish^4-7^, and 3) the adult mammalian heart maintains has a low level of proliferative cardiomyocytes maintained in the adult myocardium^29^. An additional hypothesis that has gained momentum is that cardiomyocyte proliferation is rapidly loss from microenvironment changes such as increased oxidative stress, and mitigating these changes can promote cell cycle progression *in vivo*^9, 24, 25^.

Based on the proposed importance of oxidative stress on supporting cardiomyocyte renewal, we evaluated whether decellularized matrix materials can act as a ROS sink. Studies of decellularized materials have mainly highlighted their microenvironmental cues for stimulating cellular responses, migration, and differentiation. Our investigation has highlighted an additional native functionality of decellularized myocardial matrix as directly interacting with ROS and shielding cells from its detrimental effects, which is an underemphasized role in the biomaterials field. Notably, since specific amino acid residues act as sites for oxidation such as thiols groups in free cysteine residues along with histidine and methionine^30^, compositional changes in the ECM and related changes in amino acid residue content should influence the degree this ROS scavenging activity can occur. Specifically, collagens, which are known to mainly contain glycine-proline-and hydroxyproline residue repeats, would provide minimal ROS scavenger activity compared to other ECM proteins, and ECM turnover responses that increase collagen content such as with aging^10^ and fibrosis, would be expected to reduce this capability. In comparison to the native antioxidant content, 380 ± 48 mg reduced glutathione/g tissue or 1.24 ± 0.16 µmol reduced glutathione/mg tissue has been reported for healthy porcine myocardium ^31^. Therefore, the measured value of 75.77 ± 13.16 mmol reduced glutathione equivalent/mg from our myocardial matrix material contributes a high concentration of free thiols that can greatly exceed the ROS scavenging capability found in the native myocardium alone. This is particularly evident in our *in vitro* oxidative stress assay, and previously reported results^22^, where encapsulated cardiomyocytes were shielded from the oxidative stress from H_2_O_2_ at millimolar concentrations while micromolar concentrations are able to induce extensive cell death for standard 2D cardiomyocytes cultures^32^.

For our in vivo assessments, we utilized a serial delivery of thymidine analogs to track incidents of DNA synthesis over two timespans in a single animal. Based on our assessments, the myocardial matrix hydrogel promoted around a two-fold increase in thymidine analog incorporating cardiomyocyte nuclei. This number is modest compared to some previous studies. However, it should be noted that studies reporting higher incidents have commonly utilized a cytoplasmic stain such as Troponin T or α-actinin^33^, which has risks of determining false positives due to immune cellular infiltration and potential for misidentifying overlapping cellular nuclei in the injured cardiac tissue. In comparison to these studies, cellular densities within a more conservative range from colocalized thymidine analog incorporation or positive staining for cell cycling markers have also been reported with use of PCM-1^34^. Other regenerative therapeutics such as cellular or genetic therapies have led to a large number of new cardiomyocytes in the injured myocardium by directly delivering exogenous cardiomyocyte populations or activate cardiomyocyte cell cycle progression, respectively, *in vivo*. However, safety risk of developing arrythmia from limited functional engraftment of exogenously delivered stem cells^35^ or excess promotion of endogenous cardiomyocyte turnover^36^ have limited translational potential of these direct methods for restoring *in vivo* cardiomyocytes post-MI. Injectable myocardial matrix hydrogels have been extensively evaluated for safety in animal models^15, 37^ and human patients^17^, thus, providing a safe platform for stimulating these types of endogenous repair mechanisms *in vivo*.

## Conclusion

Results from this study provides further evidence that the myocardial matrix material provides a proliferative microenvironment supporting mild cardiomyocyte renewal. As decellularized materials have been demonstrated to stimulate a plethora of endogenous repair mechanisms^36^, more mild stimulation of endogenous repair mechanisms across various cell populations instead of directly targeting a therapeutic single cellular response might better mimic native physiological repair leading to distinct improved tissue outcomes and reduced safety risks^15^. Furthermore, based on literature supporting the importance of mitigating oxidative stress for promoting cardiomyocyte renewal, decellularized materials were evaluated for directly interacting with ROS and determined in this study to act an effective ROS scavenger. Benefits of the material property as an ROS scavenger is not only relevant for cardiomyocytes, but also other cell types and various tissue diseases that oxidative stress negatively influences^37, 38^, thus, highlighting the versatility of these decellularized material therapeutics for treating chronic tissue diseases. Future work from these results could investigate the ROS scavenger activity relevant to other tissue diseases or could further enhance the ROS scavenger activity of these types of material. As injectable hydrogels have shown extensive capability as a delivery vehicle^42, 43^ or a platform for further material modifications^44^, various engineering techniques are available to further enhance therapeutic effect based on the principles highlighted in this study.

## Acknowledgements

The authors would like to acknowledge Jesus Olvera and Cody Fine of the UC San Diego Human Embryonic Stem Cell Core Facility for technical assistance of flow cytometry experiments with all made possible in part by the CIRM Major Facilities grant (FA1-00607) to the Sanford Consortium for Regenerative Medicine. This work was supported by the NIH NHLBI (R01HL113468, R01HL146147). RMW was supported through the NHLBI as a T32 training grant recipient (2T32HL105373-06A1) and a NIH F31 Predoctoral fellowship (F31HL137347). KLC is co-founder, consultant, board member, and holds equity interest in Ventrix, Inc.

## Supplementary Figures

**Figure S1:**
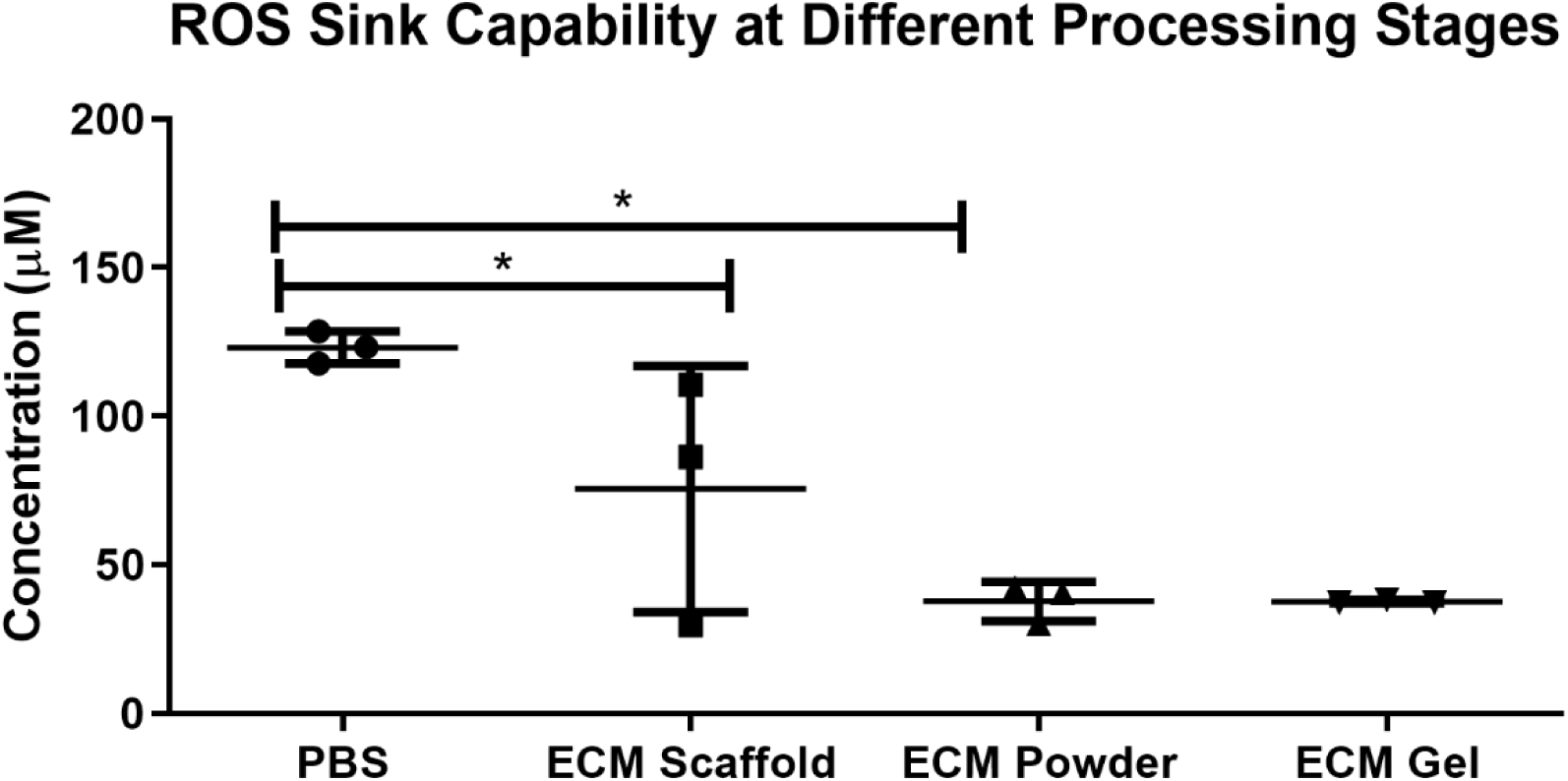
ROS sink capability at different material processing stages. 150 µL of 166.7 µM hydrogen peroxide diluted in PBS solution was applied to a solution only PBS control, a piece of lyophilized decellularized myocardial ECM, ECM powder after milling, and a myocardial matrix hydrogel (n = 3 each).

**Figure S2:**
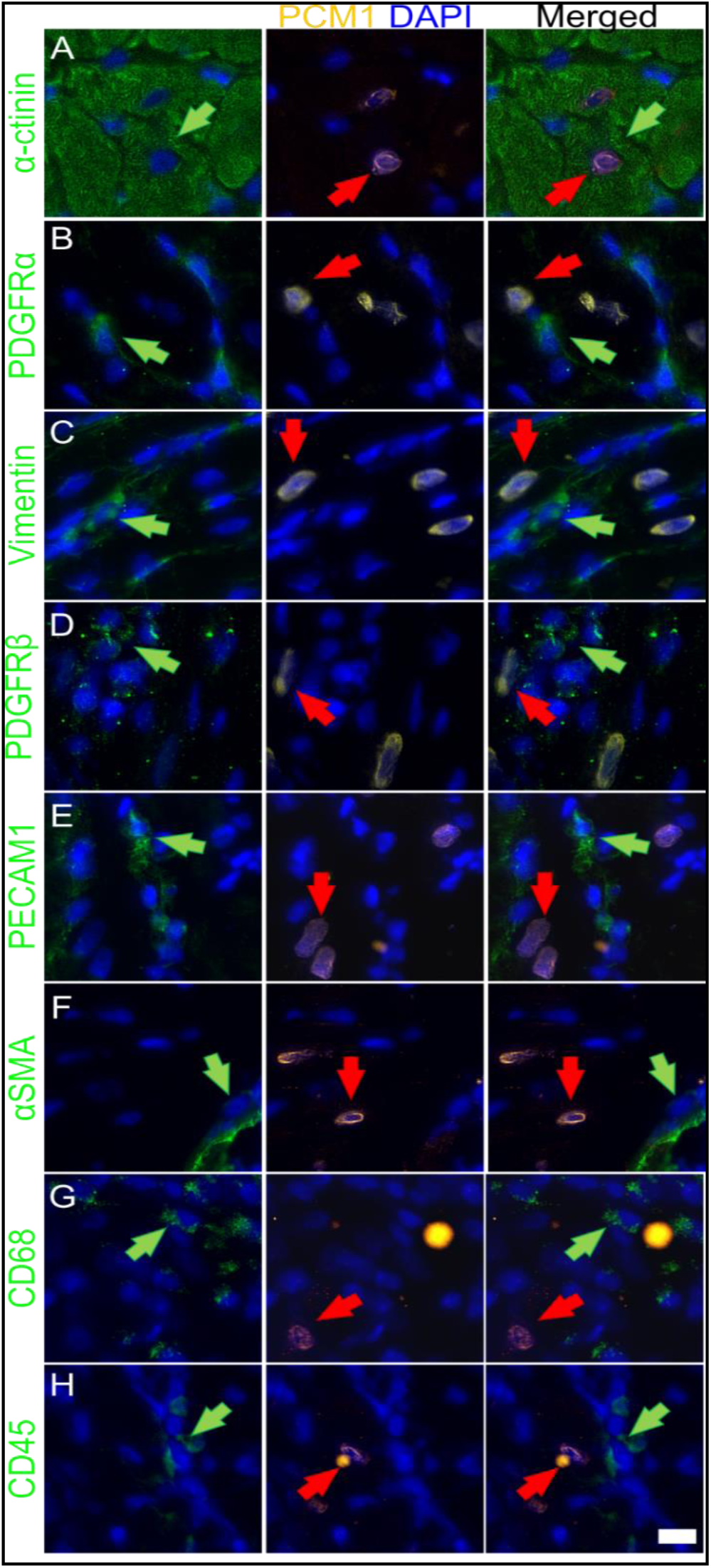
Confirmation of PCM-1 as a cardiomyocyte nuclei specific marker. (A-H) Representative images showing PCM-1 labeling overlaps with a cardiomyocyte marker (A) and does not colocalize with other cardiac cell type markers (B-H). Left column shows staining with the marker listed on the left (green). Middle column shows PCM-1 staining (orange). Right column shows merged image. Nuclei staining (blue) is present in all images. Positive staining for cardiac cell markers is indicated with a green arrow and PCM-1 with a red arrow. Scale bar is 10 µm.

**Figure S3.**
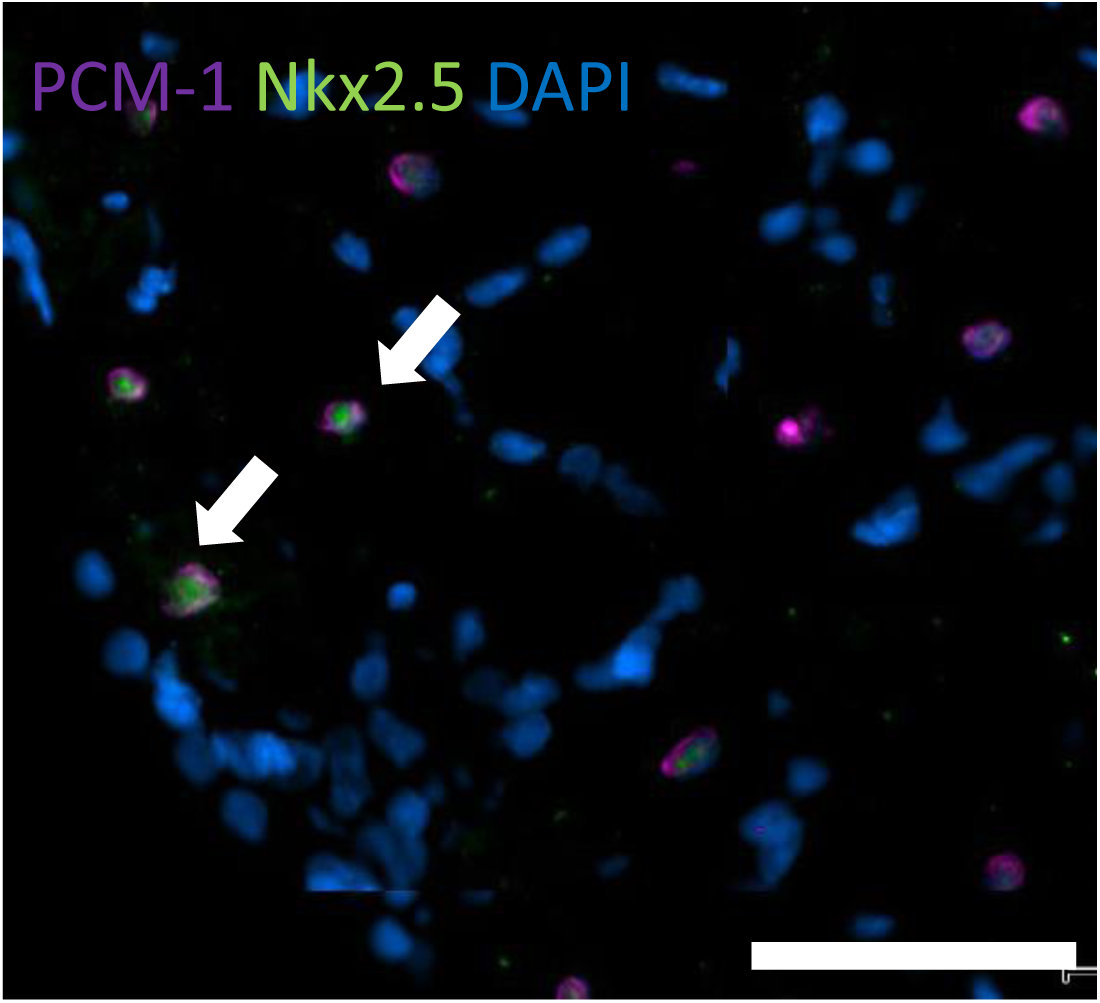
Colocalization of PCM-1 with another cardiomyocyte nuclei marker. Representative image of costaining of PCM-1 and Nkx2.5 in cardiac tissue. PCM-1 (magenta) and Nkx2.5 (green) were stained in infarcted heart tissue samples demonstrating colocalization (white arrow) that was observed throughout the tissue. Scale bar is 50 µm.

**Figure S4:**
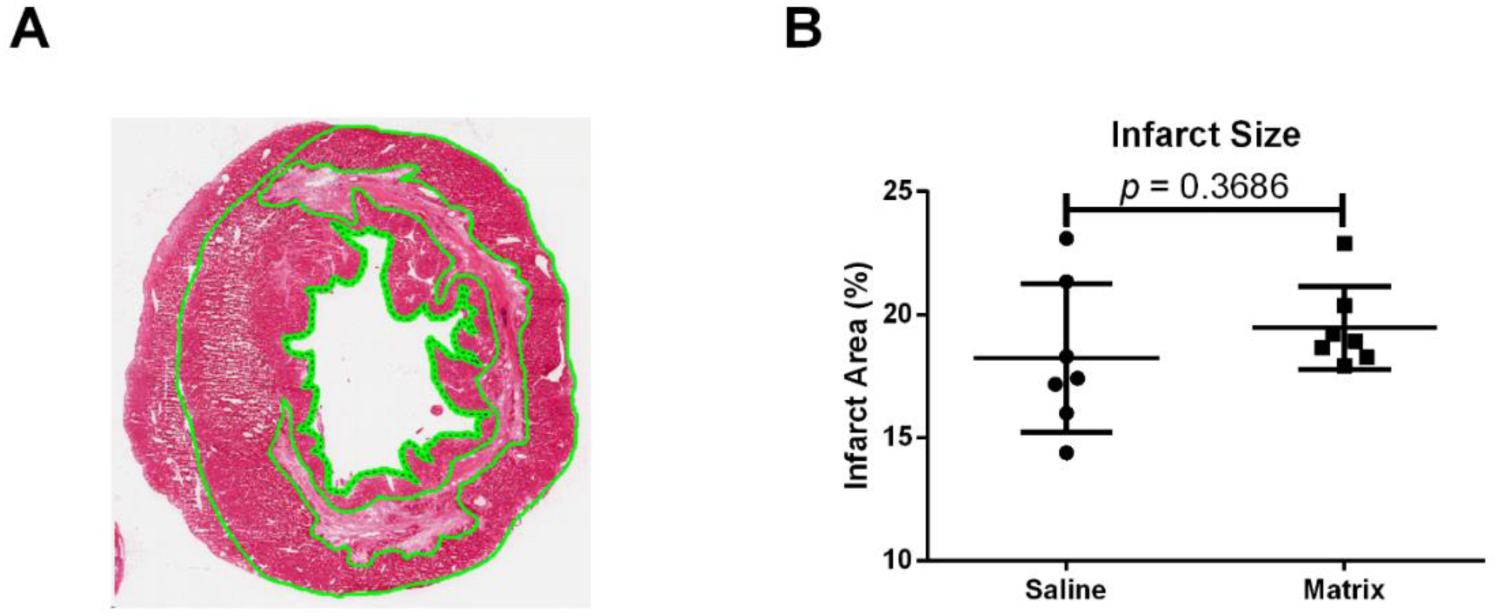
Percent infarct size analysis. (A) Image of analyzed heart for percent infarcts by highlighting (green) of the infarcted area, tissue area including left ventricle and septum, and the lumen area. (B) Quantified percent infarct size of the matrix versus saline treated hearts.

